# Investigating the human host - ssRNA virus interaction landscape using the SMEAGOL toolbox

**DOI:** 10.1101/2021.12.02.470930

**Authors:** Avantika Lal, Mariana Galvao Ferrarini, Andreas J. Gruber

## Abstract

Viruses are intracellular parasites that need their host cell to reproduce. Consequently, they have evolved numerous mechanisms to exploit the molecular machinery of their host cells, including the broad spectrum of host RNA-binding proteins (RBPs). However, the RBP interactome of viral genomes and the consequences of these interactions for infection are still to be mapped for most RNA viruses. To facilitate these efforts we have developed SMEAGOL, a fast and user-friendly toolbox to analyze the enrichment or depletion of RBP binding motifs across RNA sequences (https://github.com/gruber-sciencelab/SMEAGOL). To shed light on the interaction landscape of RNA viruses with human host cell RBPs at a large scale, we applied SMEAGOL to 197 single-stranded RNA (ssRNA) viral genome sequences. We find that the majority of ssRNA virus genomes are significantly enriched or depleted in binding motifs for human RBPs, suggesting selection pressure on these interactions. Our analysis provides an overview of potential virus - RBP interactions, covering the majority of ssRNA viral genomes fully sequenced to date, and represents a rich resource for studying host interactions vital to the virulence of ssRNA viruses. Our resource and the SMEAGOL toolbox will support future studies of virus / host interactions, ultimately feeding into better treatments.

## Introduction

According to Baltimore’s classification, class IV and class V viruses have single-stranded RNA (ssRNA) genomes ^1^. Whereas (+)ssRNA class IV viruses package the positive-sense genome that can be directly translated into protein by the translational machinery of the host cell, the (-)ssRNA class V viruses contain a negative-sense genome that needs to be transcribed into a positive-sense message before translation. ssRNA viruses interact with many host factors in the infected cells in order to facilitate viral replication, subgenomic RNA transcription, and translation of viral proteins. At the same time, host cellular factors detect viral RNA and activate intracellular signaling pathways leading to antiviral responses. Interactions between viral RNAs and host RNA-binding proteins (RBPs) are key to these processes.

ssRNA viruses such as the Hepatitis C virus (HCV), the Ebola virus, the Influenza virus, and the SARS-CoV-2 virus responsible for the ongoing COVID-19 pandemic are of high epidemiologic relevance. Understanding how these viruses interact with and impact host cells is key for designing means to combat these infections. A currently prominent example is the SARS-CoV-2 genome, which is bound by hundreds of human proteins ^2,3^. More broadly, coronaviruses are known to co-opt human RBPs to promote their stability, translation and replication ^4^. Furthermore, viral RNAs may also sequester RBPs to influence gene expression in the host. For instance, the Sindbis virus was found to “sponge” ELAVL1 RBP molecules via uridine (U)-rich elements (UREs) in its 3’ untranslated region (UTR) causing changes in splicing, polyadenylation and stability of host messenger RNAs (mRNAs) ^5^. Although studies on RBPs and viral genomes point to the importance of RBP interaction networks in viral infections, genome-scale experimental and functional studies are relatively sparse and are cell type and condition specific.

In order to bridge this gap using computational predictions, we have developed SMEAGOL (Sequence Motif Enrichment And Genome annOtation Library), a python library to analyze RBP binding motifs in nucleic acid sequences. SMEAGOL further identifies proteins whose binding motifs are significantly enriched or depleted in a sequence, thus highlighting the interactions that are most likely under evolutionary selection and therefore functionally significant. By applying SMEAGOL to 197 Group IV and Group V viral genomes we have constructed the first comprehensive resource for studying ssRNA virus/RBP interactions.

## Results

### Identification of sequence motif enrichment / depletion using SMEAGOL

SMEAGOL (https://github.com/gruber-sciencelab/SMEAGOL) is a Python library designed for comprehensive motif occurrence analysis in nucleic acid sequences using position weight matrices (PWMs), which can represent the binding specificity of a variety of nucleic acid-interacting regulators. SMEAGOL can directly load PWMs from the ATtRACT and RBPDB databases of RBP binding specificities ^6,7^. As curated databases of RBP binding motifs typically contain a mixture of high and low confidence PWMs, SMEAGOL also includes modules to analyze, filter, compare, cluster, and visualize PWMs. Moreover, SMEAGOL enables scanning of sequences with the curated PWMs and to filter these results. Post-processing modules enable the calculation and visualization of statistical enrichment or depletion of sequence motifs, as well as predicted effects of sequence variants on PWM sites. An overview of the SMEAGOL functionalities is provided in Fig. 1.

**Fig. 1.**
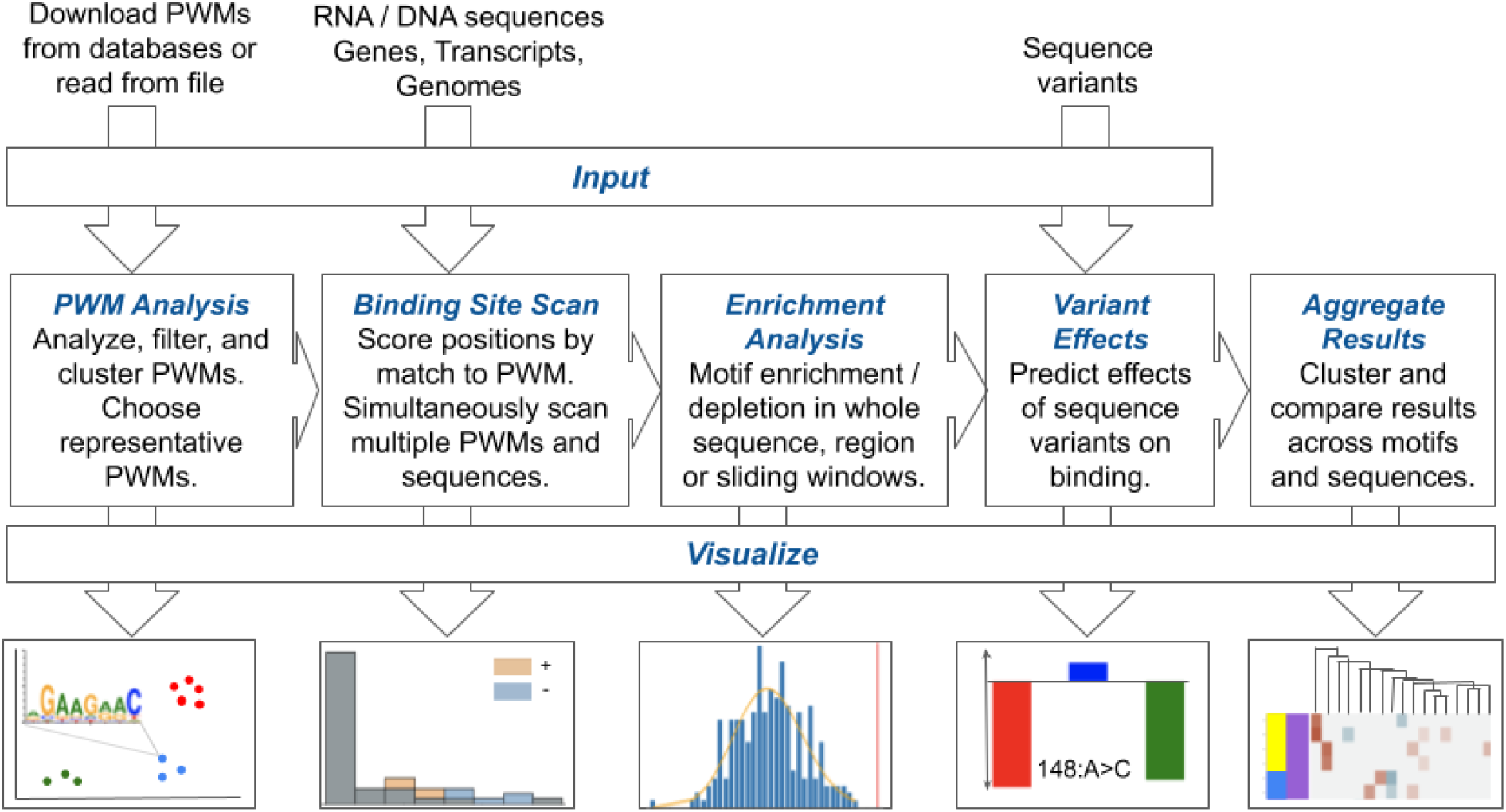
SMEAGOL enables the investigation of regulatory binding motif enrichment and variant effects in viral genomes. SMEAGOL takes as input sequence files in FASTA format and regulator binding specificities in the form of position weight matrices (PWMs) in order to perform PWM analysis, sequence scanning, enrichment / depletion analysis, and variant effect prediction. Finally, SMEAGOL enables visualization of the results in various ways.

### The genomes of ssRNA viruses show evidence of selection on RBP binding sites

In order to find out which RBPs are enriched / depleted in binding sites across ssRNA virus genomes, we have used PWMs representing the experimentally determined sequence binding preferences of human RBPs from the ATtRACT ^7^, RBPDB ^6^, and ENCODE ^8^ databases. We used SMEAGOL to filter and curate these motifs (Methods, Supplementary Fig. 1) and obtained a curated set of 362 PWMs representing the binding specificities of 146 human RBPs. We then used this set of PWMs and SMEAGOL to scan the complete genome sequences of 197 ssRNA viruses belonging to 19 families (Table 1, Supplementary Data 1) in order to identify putative RBP binding sites (Supplementary Data 2).

**Table 1:** Viral classes and families as well as number of species and genomes considered within this study.

| Baltimore Classification | Type | Genome Info | Family | Number of Genomes | Number of Species |
| --- | --- | --- | --- | --- | --- |
| Class IV | (+)ssRNA | Segmented | *Flaviviridae* * <sup>1</sup> | 2 | 2 |
|  |  | Monopartite | *Astroviridae* | 11 | 11 |
|  |  |  | *Caliciviridae* | 2 | 2 |
|  |  |  | *Coronaviridae* | 8 | 8 |
|  |  |  | *Flaviviridae* * <sup>1</sup> | 37 | 31 |
|  |  |  | *Hepeviridae* | 1 | 1 |
|  |  |  | *Matonaviridae* | 1 | 1 |
|  |  |  | *Picornaviridae* | 22 | 21 |
|  |  |  | *Togaviridae* | 11 | 11 |
|  |  | Total |  | 95 | 88 |
| Class V | (-)ssRNA | Segmented | *Arenaviridae* | 9 | 9 |
|  |  |  | *Hantaviridae* | 9 | 9 |
|  |  |  | *Nairoviridae* | 3 | 3 |
|  |  |  | *Orthomyxoviridae* | 10 | 4 |
|  |  |  | *Peribunyaviridae* | 22 | 22 |
|  |  |  | *Phenuiviridae* | 10 | 10 |
|  |  | Monopartite | *Bornaviridae* | 3 | 2 |
|  |  |  | *Filoviridae* | 7 | 6 |
|  |  |  | *Paramyxoviridae* | 15 | 15 |
|  |  |  | *Pneumoviridae* | 2 | 2 |
|  |  |  | *Rhabdoviridae* | 12 | 12 |
|  |  | Total |  | 102 | 94 |

\*1: Two genomes included in the *Flaviviridae* family are evolutionarily related to the monopartite viruses of the genus *Flavivirus*, but belong to a group of segmented, and likely multicomponent viruses, the jingmenviruses<sup>9</sup>.

Finally, to find evidence of evolutionary selection for or against RBP binding to viral RNA we determined the enrichment or depletion of RBP binding motifs in each viral genome compared to dinucleotide-randomized versions of the genome (Supplementary Data 3). It should be noted that, although the viruses in our study have single-stranded genomes, the complementary strand of the genome is also synthesized during the viral life cycle and interacts with host factors. We therefore repeated this procedure for the complementary strand of each genome.

Overall, we found that (+)ssRNA (Class IV) virus genomes had more motif enrichment or depletion in both strands compared to (-)ssRNA (Class V) virus genomes (Supplementary Fig. 2). However, the number of motifs enriched or depleted also varied significantly between families within each group (Fig. 2a).

**Fig. 2.**
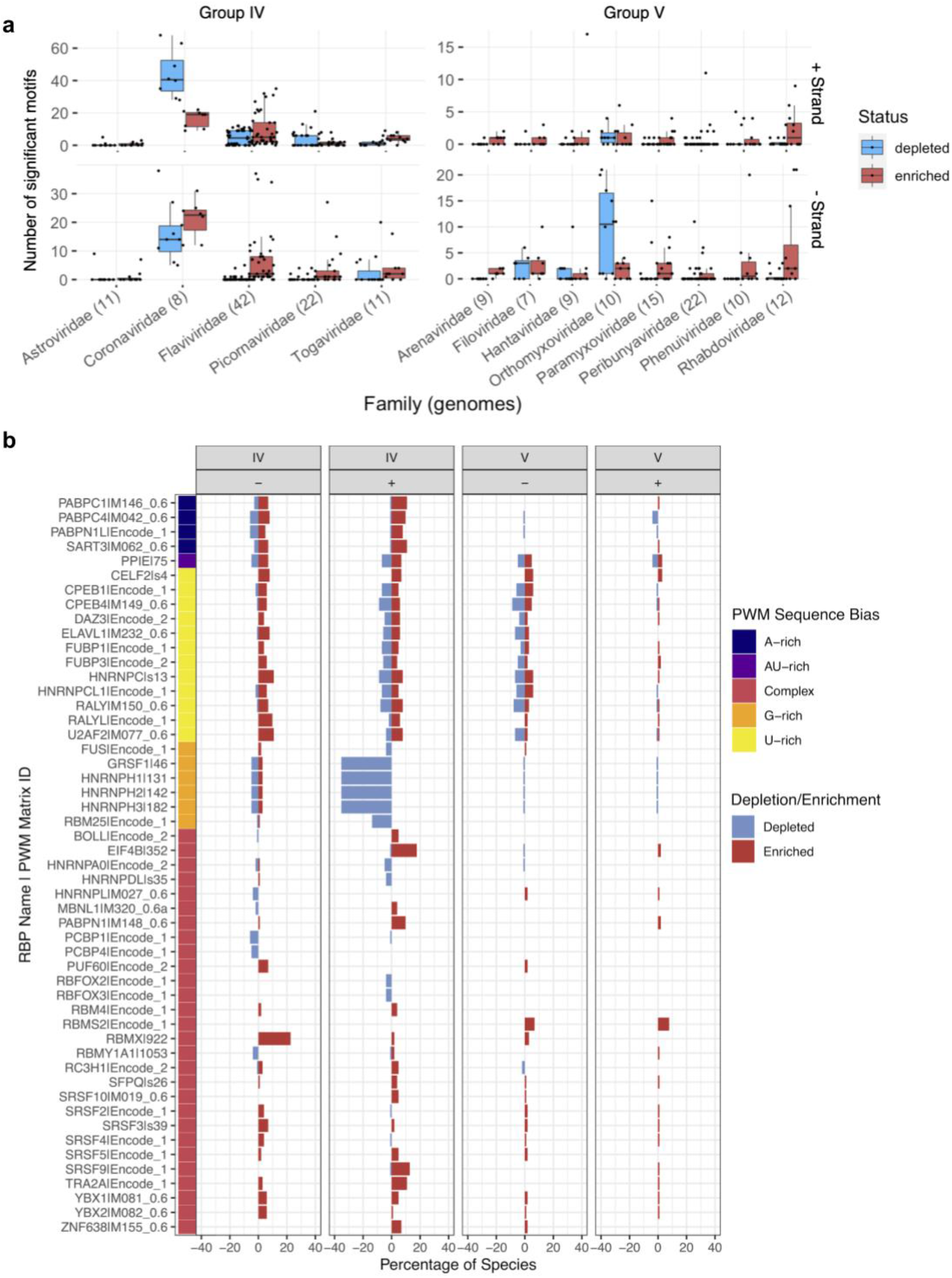
SMEAGOL uncovers RBPs whose binding motifs are enriched or depleted in ssRNA virus genomes. **a** | Number of motifs significantly enriched or depleted in each viral family. The number of genomes per family is given in parentheses. Families with more than 5 genomes in our dataset were included. Box plots are defined as follows: center line, median; box limits, upper and lower quartiles; whiskers, 1.5x interquartile range. Individual data points are also shown. **b** | Percentage of viral genomes with significant (two-sided binomial test, FDR-adjusted p-values < 0.05) enrichment (in red) and depletion (in blue) per PWM, separated by virus class (IV or V) and viral genome strand. For readability, shown are only cluster-representative PWMs that had more than three significant enrichment / depletion events in at least one of the populations (group IV or V in strand + or -). PWM sequence bias (see Methods) is presented on the left side of the plot. While some PWMs have a more complex sequence (in pink), others are rich in single nucleotides (A-rich in navy blue, AU-rich in purple, U-rich in yellow, and G-rich in orange). A complete figure containing all cluster-representative PWMs is provided as Supplementary Fig. 3.

Interestingly, while some RBP binding motifs are generally depleted or enriched in (+)ssRNA virus genomes, other motifs are much more specific to a subset of viruses. For instance, G-rich motifs recognized by several splicing factors (GRSF1, HNRNPH1, HNRNPH2, HNRNPH3, HNRNPF, HNRNPA2B1) are frequently depleted in the plus-strand of (+)ssRNA virus genomes across multiple families (Fig. 2b and Supplementary Data 3).

On the other hand, the motif for RBMX is enriched on the negative sense molecule in several families of both groups IV and V (28 genomes from four families in our dataset). All four Dengue virus (DENV) genomes in our dataset showed enrichment in the negative sense molecule for RBMX (Supplementary Data 2). Interestingly, RBMX is required for efficient amplification of DENV and its knockdown significantly decreased DENV titer ^10^. The second most commonly enriched motif was related to the translation factor EIF4B, specifically in the positive strand of (+)ssRNA viruses. EIF4B was reported to bind to DENV RNA ^11^ and its depletion reduced the efficiency of translation initiation in zika virus (ZIKV) ^12^. All DENV and ZIKV genomes in our dataset had significant enrichment for this host factor as did most flaviviruses (Fig. 3).

**Fig. 3.**
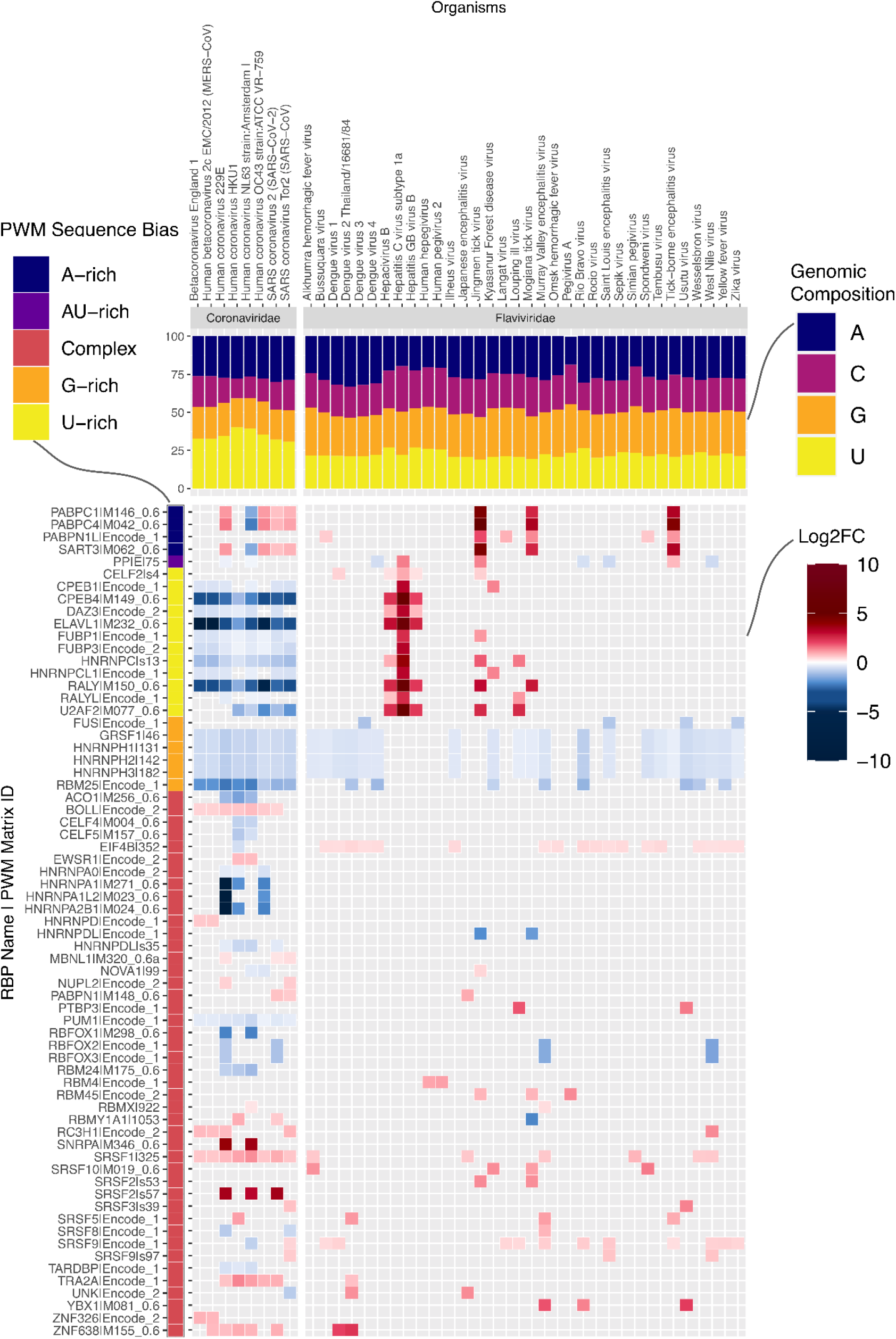
Enrichment and depletion results for *Coronaviridae* and *Flaviviridae* families. Heatmap with results for positive sense molecules in terms of log2 fold-change (log2FC) of enrichment (in red hues) or depletion (in blue hues) for single genomes within two (+)ssRNA families. For reasons of space, only cluster-representative PWMs enriched or depleted in more than one viral genome are shown (species-specific results are provided in Supplementary Fig. 6). The PWM sequence bias is presented on the left side of the plot (as explained in Fig. 2). The nucleotide compositions of the viral genomes are provided on the top of the heatmap (A in navy-blue, C in magenta, G in orange, U in yellow).

Having observed significant differences between viral families, we next examined prominent families individually. Coronaviruses, which have the longest genomes of all viruses in our dataset (Supplementary Fig. 4, one-sided Wilcoxon rank-sum test U statistic = 1536, p = 8.5 × 10^−7^), also show more enrichment and depletion of binding motifs than any other family (Fig. 2a). Strikingly, the number of RBP motifs depleted on the plus strand is much higher than the number of enriched motifs. It is conceivable that given their long genomes, coronaviruses have to actively prevent being bound by non beneficial host RBPs. On examination, we found that the striking number of depleted motifs in these genomes reflects depletion of U-rich elements bound by RBPs such as HNRNPC, RALY, CELF2, TIA1, ELAVL1 and PPIE (Fig. 3). This is despite coronaviruses being the most U-rich of all viruses in our dataset (Fig. 3, Supplementary Fig. 5, one-sided Wilcoxon rank-sum test U statistic = 1530, p = 5.4 × 10^−13^). By contrast, a subset of flaviviruses, which are relatively poor in uridines, are enriched for these UREs (Fig. 3, Supplementary Fig. 5). In addition to the depletion of UREs and G-rich elements, all 8 coronaviruses in our dataset showed enrichment of motifs for the SRSF1 splicing factor and depletion for the PUM1 RBP, which reduces mRNA stability ^13^.

**Fig. 4.**
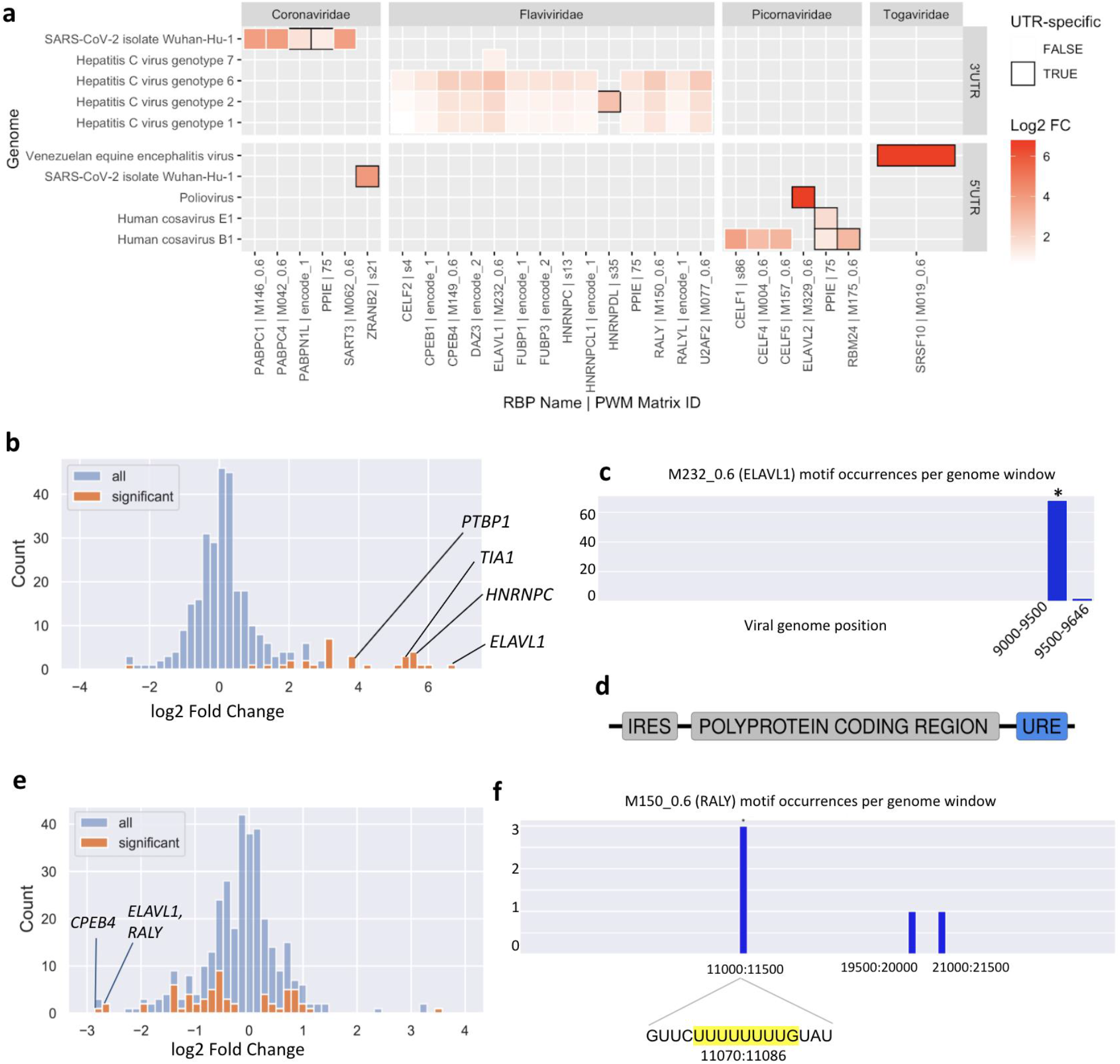
Motif enrichment patterns in specific regions of viral genomes. **a** | UTR enrichment / depletion analysis results from SMEAGOL. Cells with black frames represent motifs that were not significantly enriched in the whole genome. Only cluster-representative motifs (see Methods) were included in this figure. Full results are given in Supplementary Data 3. **b** | Motifs for 22 RBPs were predicted by SMEAGOL to be significantly enriched within the HCV genome while 1 motif was depleted (shown are the results for HCV genotype 1a). **c** | Using the local window enrichment function of SMEAGOL (see Methods), we found that the vast majority of regions lack binding motifs for the ELAVL1 (HuR) RBP, whereas it was highly enriched within a region at the 3’ end of the virus genome (Two-sided Fisher’s exact test, odds = 21.8, FDR-adjusted p-value = 1.4 × 10^−51^). **d** | A U-rich region is located within the 3’ UTR of the HCV genome. **e** | RBPs predicted by SMEAGOL to be significantly enriched / depleted in binding sites within the SARS-CoV-2 genome. URE-binding RBPs are shown to be most strongly depleted. **f** | Using the local window enrichment function of SMEAGOL, we observed that binding motifs of RALY are absent in most of the SARS-CoV-2 genome but enriched in a specific region (Two-sided Fisher’s exact test, odds = 36.5, FDR-adjusted p-value = 0.013). On closer inspection, this is due to multiple motifs within an URE at position 11074.

### Non-coding regions in viral genomes show distinct patterns of enrichment

Like cellular mRNAs, the genomes of (+)ssRNA viruses also contain 5’ and 3’ UTRs, which have been shown to bind host RBPs. While the 5’ UTR contains elements that regulate the efficiency and timing of translation initiation and viral replication, host factors binding to the 3’ UTR can be critical to many aspects of the life cycle of a virus, including but not limited to RNA replication and stability. Host RBPs also mediate 5’ UTR - 3’ UTR interactions, resulting in ‘circularization’ of the viral genome ^14,15^.

Since the UTR regions have distinct regulatory functions from the remaining genome and their sequences are not constrained to code for proteins, we reasoned that they may be enriched for binding sites of specific RBPs relevant to their functions. These enrichments may not be detectable over the whole genome, and indeed may be canceled out since it may be detrimental for some UTR-specific proteins to bind elsewhere in the genome. We therefore repeated the analysis specifically for 5’ and 3’ UTR sequences of 89 (+)ssRNA viruses whose UTR positions were annotated.

This analysis highlighted new putative host-virus associations. Of 116 enriched motif-UTR pairs (Supplementary Data 4), 22 were not enriched in the whole genome of the same virus. In general, 3’ UTRs exhibited more instances of motif enrichment compared to the 5’ UTR sequences. One consistent result was an enrichment of U-rich motifs in the 3’ UTRs of multiple Hepatitis C Virus (HCV) genotypes (Fig. 4a).

To further validate the predictions made by SMEAGOL, we focused on two well-studied pathogenic (+)ssRNA viruses, namely HCV, from the Flaviviridae family, and SARS-CoV-2, from the Coronaviridae family. The interactions of these viruses with host factors have been studied experimentally and both are enriched / depleted in binding sites of specific RBPs according to our analysis.

### The Hepatitis C virus genome is highly enriched in binding sites of U-rich element binding RBPs

Among the top ten significant RBPs whose motifs are enriched / depleted in the HCV genome, four interactions have already been experimentally validated. The RBP that is most significantly enriched in binding sites in the HCV genome is the ELAV-like RNA binding protein 1 (ELAVL1), also called HuR (Fig. 4b). Multiple experimental studies have reported direct binding of ELAVL1 to the HCV genome ^16–18^ and RNAi experiments have shown that ELAVL1 knockdown counteracts HCV replication ^17,19,20^. ELAVL1 activates translation mediated by the HCV internal ribosome entry site (IRES) via an unknown mechanism which seems to not require direct binding of ELAVL1 to the IRES itself ^21^. Using the local enrichment function of SMEAGOL we found that the HCV genome is highly enriched in ELAVL1 motifs within its 3’ UTR (Fig. 4c), consistent with previous reports that ELAVL1 directly interacts with a U-rich region located within the 3’ UTR of the virus ^16,17^ (Fig. 4d).

On the role of the RBP for the reproduction of the virus, ELAVL1 was reported to displace the Polypyrimidine Tract Binding Protein 1 (PTBP1) to facilitate binding of the La Protein (SSB) to the HCV 3’ UTR and enhance viral replication ^22^. Strikingly, PTBP1 was also one of the most enriched RBPs predicted by SMEAGOL. PTBP1 was reported to bind the HCV genome by several studies ^23–30^. Specifically, it interacts with the 5’ UTR as well as the 3’ UTR ^27^, although binding to the 5’ UTR is much weaker than to the 3’ UTR ^25^. PTBP1 silencing blocks HCV replication ^31,32^ and decreases the amount of viral RNA ^20^. In mice, PTBP1 was found to play a role in HCV transcription ^33^. The RBP can act in the nucleus as well as in the cytoplasm ^34^ and was found to colocalize with the HCV NS5B protein ^32^.

Another highly enriched RBP that was previously reported to directly interact with the virus was the Heterogeneous Nuclear Ribonucleoprotein C (HNRNPC). Multiple studies have shown that HNRNPC physically interacts with the HCV genome ^16,17,24^. The RNA recognition motif (RRM) of HNRNPC binds to tracts of four or more uridines ^35–37^, with higher affinity for longer tracts ^37^. Accordingly, HNRNPC binds to the poly(U) tract in the HCV 3’ UTR ^24,38^. Interestingly, an RNAi study has shown that *HNRNPC* knockdown decreases cellular HCV RNA levels ^20^ which indicates that *HNRNPC* positively contributes to HCV replication.

Finally, the TIA1 RBP was also among the most enriched RBPs predicted by SMEAGOL. HCV infection causes autophosphorylation of protein kinase R (PKR) thereby repressing the translation of capped cellular mRNAs ^39^ and forcing the formation of stress granules (SGs) ^40^. Importantly, it was reported that the stress granule associated protein TIA1 is required for efficient HCV production at early and late time points of infection. This suggests that HCV exploits the SG machinery of its host for efficient reproduction. However, the study also demonstrates that TIA1 is required for efficient HCV infection even in the absence of SG formation ^41^. The viral NS5A protein colocalizes with TIA1 in association with lipid droplets, which are important organelles of viral assembly, and is immunoprecipitated using antibodies to TIA1. In a study that combined RNA affinity capture with mass spectrometry, TIA1 has been found to associate with the HCV 3’ UTR ^17^. Also, TIA1 knockdown was reported to reduce HCV RNA and infectivity in Huh-7 cells ^41^.

The remaining six RBPs whose motifs are most significantly enriched in the HCV genome are RALY, CPEB4, HNRNPCL1, U2AF2, TRNAU1AP and BOLL. Further down the list, we can see more interesting candidates enriched in the HCV genome, including FUBP1 and YBX1. FUBP1 was previously reported to directly interact with the viral 3’ UTR ^17^ and also appears to facilitate persistent replication of HCV by regulating p53 ^42^. Similarly, there exists experimental evidence for direct interaction of YBX1 with the HCV genome ^17,43^.

### Motif enrichment expands upon functional studies in the SARS-CoV-2 genome

The enrichment and depletion of RBP motifs in the SARS-CoV-2 genome is largely similar to that of the other coronaviruses in our dataset. However, it is unique in having strong enrichment for binding motifs of YBX1, which has been experimentally validated to bind to SARS-CoV-2 RNA ^2^ and supports infection by other viruses, including Dengue Virus ^44^, Influenza ^45^ and HIV ^46^. Like other coronaviruses, more RBPs were enriched and depleted (18 and 32 respectively) in the positive sense genome sequence compared to the negative sense intermediates (14 and 13 respectively), suggesting that more functional interactions happen with the positive sense molecule. In the genome, we found motif depletion to be more common than enrichment (motifs for 32 RBPs were depleted while motifs for 18 RBPs were enriched), suggesting that SARS-CoV-2 has more antiviral interactions with human RBPs than pro-viral interactions. This prediction is consistent with experimental observations from CRISPR screens ^2^.

To place our predictions for specific RBPs in the context of experimental data, we collected a list of proteins that have been experimentally validated to bind to SARS-CoV-2 RNA in infected human or monkey cell lines in three studies ^3,47,48^. PWMs for 41 of these were included in our study, and we computationally predicted binding sites for 40 of these 41 in the SARS-CoV-2 genome. We found motifs for 8 of these RBPs to be enriched while 13 were depleted, indicating that while some interacting RBPs bind to longer regions or an abundance of locations in the viral genome, others are overall depleted in binding sites in order to guarantee highly specific binding to well defined genomic loci.

We also compiled a list of experimentally validated antiviral and pro-viral RBPs from CRISPR or siRNA screens in SARS-CoV-2 infected cells ^47,49^. PWMs for 17 known antiviral and 4 known pro-viral RBPs were included in our dataset. While we did not observe motif enrichment for the pro-viral proteins, motifs for 4 of the 17 antiviral proteins (RALY, ELAVL1, FUBP3, PCBP2) were depleted in the SARS-CoV-2 genome, suggesting that the viral genome may have evolved to avoid interaction with these defensive host proteins. Motifs for an additional 4 antiviral RBPs (HNRNPA2B1, DAZAP1, TARDBP, PPIE) were also depleted at a more permissive FDR-adjusted p-value threshold of 0.1. Out of these, RALY, ELAVL1, FUBP3 and PPIE bind to UREs. Interestingly, although the predicted binding sites for numerous URE-binding RBPs are strongly depleted overall in the SARS-CoV-2 genome (Fig. 4e), the few binding sites that are predicted are significantly concentrated within a region in the NSP6 gene. In particular, an URE at position 11074 contains 3 of 5 predicted binding sites for the antiviral RBPs RALY and ELAVL1 (Fig. 4f).

Computational studies offer an opportunity to predict novel interactions that may not have been covered in the limited range of cell types and conditions that were studied experimentally. We identified strong (FDR-adjusted p-value < 0.05 and fold change >= 2) enrichment of motifs for 5 RBPs (SART3, PABPC1, NUPL2, SRSF2, ZRANB2) and strong depletion (FDR-adjusted p-value < 0.05 and fold change <= 0.5) of motifs for 10 RBPs (CPEB4, HNRNPA1, HNRNPC, HNRNPCL1, HNRNPK, RBFOX1, RBFOX2, RBFOX3, RBM25, U2AF2) that were not listed as hits in the screens we examined.

### SMEAGOL offers functionality to predict sequence mutation effects on RBP binding

Since computational analysis allows us to predict the probable locations of protein binding on the viral genome, it also offers the possibility of predicting how mutations may affect these binding sites. To demonstrate this functionality, we selected 10 RBPs that (1) have PWMs in our dataset, (2) are experimentally determined to be antiviral in SARS-CoV-2 infection and/or are shown by our analysis to be significantly depleted in the SARS-CoV-2 genome, and (3) have fewer than 10 predicted binding sites on the SARS-CoV-2 genome. These are CPEB4, RALY, ELAVL1, HNRNPA1, RBFOX1, HNRNPK, HNRNPA2B1, DAZAP1, SRSF7 and PCBP2. We hypothesized that mutations that disrupt the binding sites of these RBPs may enable SARS-CoV-2 to escape host antiviral defenses. Using SMEAGOL to analyze a database of SARS-CoV-2 mutations ^50^, we identified 170 mutations that are predicted to disrupt motifs for the selected RBPs (Supplementary Data 5). Interestingly, this list includes 22 non-exonic and 60 synonymous mutations.

As an example, the T11078C (nsp6:p.F36L) mutation, one of the lineage-determining mutations in the N.9 Variant of Interest found in Brazil ^51^, is predicted to disrupt binding of RALY, ELAVL1, and CPEB4 (Supplementary Fig. 7) to the URE at position 11074 (Fig. 4f). As discussed above, this URE is one of very few regions predicted to bind to the known antiviral RBPs RALY and ELAVL1, as well as CPEB4 which is validated to bind to the SARS-CoV-2 genome. Interestingly, the much less common T>C mutation at the same position is predicted to have a lesser effect on RBP binding (Supplementary Fig. 7). This example illustrates the capability of SMEAGOL to generate predictions for the functional effects of sequence mutations or variants, and to prioritize variants for experimental studies.

## Discussion

There exist several web servers ^52^ and libraries ^53–56^ to scan nucleic acid sequences with PWMs and identify putative binding sites. Several tools ^52,56^ also calculate a p-value for motif enrichment that takes into account the nucleotide composition of the sequence. With SMEAGOL, we aim to provide a unified python-based framework for visualization, analysis and clustering of PWMs, binding site discovery, variant effect prediction, and binding site enrichment/depletion calculations using a background model that incorporates user-specified k-mer frequency in the query sequence. For instance, dinucleotide shuffling is considered to be more conservative compared to mononucleotide shuffling, as it better accounts for RNA structural features and genomic biases in the occurrence of dinucleotides. SMEAGOL is able to perform this enrichment calculation with hundreds of PWMs on large (10 kb) sequences in under a minute (Supplementary Fig. 8). While SMEAGOL was designed with a focus on RBP-RNA interactions, it can be applied to analyze protein interactions with genomes,genomic regions, genes, or transcripts. Here, we have applied it to perform the first large-scale computational analysis of interactions between RNA viruses and human RBPs.

We found numerous RBP-binding motifs to be enriched or depleted in ssRNA viruses, including motifs that were enriched or depleted globally as well as in a family- or species-specific manner. The RBPs bound by these motifs include host splicing factors as well as RBPs that are known to regulate RNA stability. We report differences in predicted host interactions between viral families, with coronaviruses showing the highest levels of motif enrichment and depletion in their genomes. Coronaviruses may have evolved to avoid being bound by specific RBPs as, given their length, most RBPs will bind the viral genome by chance in the absence of active selection against it. Further, we find an interesting pattern in the occurrence of UREs which bind numerous RBPs that regulate viral infection. These UREs are depleted in the genomes of coronaviruses (which are highly U-rich overall) and enriched in a few flaviviruses including HCV (which are overall depleted in uridine). Consistent with our findings, the ELAVL1 RBP that binds to these elements has been experimentally found to be antiviral in SARS-CoV-2 infection but pro-viral in HCV infection.

HCV causes chronic infections of the liver, which further lead to diseases such as liver cirrhosis and hepatocellular carcinoma. About 70 million people are chronically infected by HCV and a significant number of these will develop very severe diseases, such as liver cirrhosis and malignancies ^57^. Previous studies confirm that at least four of the top ten RBPs enriched / depleted in binding sites within HCV, do indeed bind to the viral genome. These are HNRNPC, TIA1, PTBP1 and ELAVL1, all of which are URE-binding RBPs. Secretion of type I interferon (IFN) is one of the first cellular responses to viral infection. Many cytokines contain UREs in their 3’ UTRs making them unstable. Binding of ELAVL1 to UREs stabilizes transcripts. Interestingly, ELAVL1 has been shown to bind and stabilize the Interferon-beta transcript. Accordingly, ELAVL1 knockdown dampens type I IFN response ^58^. Moreover, reduced ELAVL1 expression decreased protection against Vesicular stomatitis virus (VSV) infection ^58^. These results indicate that ELAVL1 acts as a potent regulator of type I IFN response ^59^. Consequently, it is conceivable that sequestration of ELAVL1 by HCV might contribute to reduced defense against the virus due to a decreased type I IFN response. In contrast, CRISPR knockout of ELAVL1 sensitized VeroE6 cells to SARS-CoV-2 infection ^49^, indicating an antiviral effect of this RBP against SARS-CoV-2, though the mechanism is unclear.

We previously published an analysis of motif enrichment in the SARS-CoV-2 genome using a similar procedure ^60^. Here, we improve upon the previous findings with a more rigorous statistical procedure including dinucleotide shuffling, using an expanded dataset of PWMs, and by placing the results in the context of other Coronavirus genomes and more recent functional studies. Our new analysis supports the observation that SARS-CoV-2 is more likely to form antiviral interactions with RBPs than pro-viral ones. Further, we find depletion of motifs for several known antiviral RBPs on the SARS-CoV-2 genome. We extend functional studies by providing binding site predictions for known pro-viral and antiviral RBPs as well as predicting putative interactions. Finally, we predict mutations that may disrupt the binding sites of known antiviral proteins. While mutations affecting SARS-CoV-2 protein sequences have been extensively studied, the effects of other classes of mutations are less clear. SMEAGOL supports the creation of testable hypotheses and helps to prioritize non-coding and non-synonymous mutations for detailed analyses.

Our dataset of predicted RBP-virus interactions is freely available (Supplementary Data 3, Supplementary Data 4) along with our software. We suggest that the proteins highlighted in our analysis can be prioritized in knockout, knockdown or overexpression studies to experimentally measure their impact on viral pathogenesis. RBPs whose binding motifs are enriched in viral genomes may prove to be targets for antiviral drugs. These interactions may also indirectly modulate host transcription by sequestering RBPs. Follow-up studies may test this hypothesis by analyzing gene expression data for indications that human transcripts normally regulated by these RBPs are differentially expressed due to altered RNA stability or dysregulated splicing in virus-infected cells. Further, our results will foster a better understanding of factors that mediate the host species range of ssRNA viruses, the range of susceptible cell types in the human body, and the vulnerability of different human populations to viral disease.

Finally, deep learning methods have shown promise for identifying nucleic acid-protein binding sites, potentially with higher accuracy than PWM scanning ^61–63^, and tools have recently been developed to learn motif representations from the trained parameters of these models ^64^. However, trained models are not available for many human RBPs and the methods are generally difficult to use for non-experts. A possible extension of SMEAGOL in the future could be to incorporate deep learning and structure-based models, in order to offer improved predictions wherever possible and to benefit the field by enabling easier comparison between methods.

## Materials and Methods

### Curation of viral genomes

The complete genome sequences for viruses (Taxonomy ID: 10239) deposited in the NCBI repository (https://www.ncbi.nlm.nih.gov/genome/browse#!/viruses/) were retrieved using the following search / filter strategies: only RefSeq entries of specific families of (+)ssRNA and (-)ssRNA viruses known to infect humans (host = “human”) were selected (Table 1). We manually curated these data by adding missing information from additional viral databases ViPR (https://www.viprbrc.org) ^65^ and ViralZone (https://viralzone.expasy.org/) ^66^. All information regarding reference strains or the selection of representative strains/genotypes along with excluded genomes can be found in Supplementary Data 1.

The complete genomic sequences were downloaded along with the GFF annotations. For (+)ssRNA viruses, the GFF annotation was used to extract the 3’ UTR and 5’ UTR sequences wherever possible.

### Curation of PWMs

All available position matrices were downloaded from the ATtRACT database (https://attract.cnic.es/) and from the RBPDB database (http://rbpdb.ccbr.utoronto.ca/, version 1.3.1) on February 2, 2021. The matrices were filtered to retain only matrices derived from competitive binding experiments using wild-type human RBPs. RBPDB matrices which were redundant with ATtRACT were removed. PWMs from ENCODE RNA Bind-n-Seq assays ^8^ were constructed using the ENCODE computational pipeline ^8^ and added to this list. For RBPDB, position frequency matrices (PFMs) were converted to PWMs with a pseudocount of 0.01. For ATtRACT, position probability matrices (PPMs) were downloaded and converted to PWMs using the ‘ppm_to_pwm’ SMEAGOL function. 44 PWMs were trimmed using the ‘trim_ppm’ SMEAGOL function with the default parameters to remove uninformative positions. PWMs were filtered to remove outliers with an entropy greater than 10. PWMs of length 4-12 positions were retained. Ultimately, 362 PWMs representing 146 human RBPs were considered for the downstream analyses.

PWMs with high sequence bias toward one of the four bases (A, G, C, or U) were identified by scanning sequences of poly-A, poly-C, etc. PWMs that had a match score of > 0.8 to a single-base sequence were annotated as being biased toward that base.

### Selection of representative PWMs for Figures 2-4

84 RBPs had multiple PWMs in our filtered final set of PWMs. For 66 of these, we used the ‘choose_representative_mat’ function in SMEAGOL to select a single representative PWM by calculating pairwise similarities between all PWMs in the group based on the normalized Pearson correlation metric ^67^ and selecting the one which had the maximum similarity (defined as median normalized correlation) to the others. For 18 RBPs, we observed PWMs falling into dissimilar groups (specifically, the normalized correlation between at least one pair of PWMs for the RBP was below 0.2). Therefore, for each of these 18 RBPs, we used the ‘cluster_pms’ function of SMEAGOL to cluster the PWMs using agglomerative clustering based on the normalized correlation metric, and select a representative PWM for each cluster.

### Calculation of motif enrichment and depletion

The ‘scan_sequence’ function of SMEAGOL scans a given nucleic acid sequence by calculating the PWM match score for each position in the sequence. Specifically, at each position in the sequence, the subsequence of length *k* (where *k* is the length of the PWM) starting at the given position is taken, and the PWM match score is obtained by summing over the PWM log-likelihood ratios at each of the *k* positions, each time selecting the PWM element that corresponds to the nucleotide in the sequence. The score is then divided by the maximum possible score that could be obtained using that PWM ^55^. We used this function to scan the downloaded ssRNA virus genome sequences, as well as their reverse complement sequences, with the 362 selected RBP PWMs, and identified putative binding sites with a score threshold of 0.8. We used the ‘enrich_in_genome’ function in SMEAGOL to calculate a p-value for enrichment or depletion of each PWM and viral genome.

The p-value is calculated as follows. For each genome and PWM combination, SMEAGOL counts the number of predicted binding sites. It then generates 1000 background sequences that have the same nucleotide and dinucleotide frequency as the genome, scans each background sequence, and counts the number of predicted binding sites in the background sequences to generate a background distribution. This is used to calculate the expected probability of finding a binding site in the query sequence based on its sequence composition alone. A two-sided binomial test is used to calculate the p-value, which is then adjusted for multiple-testing using the Benjamini-Hochberg correction. For multi-segmented viral genomes, SMEAGOL calculates a single enrichment score across all segments.

PWMs with FDR-adjusted p-value < 0.05 were considered to be significantly enriched / depleted. The ratio of the real and expected number of binding sites in the query sequence was used as a measure of effect size.

### Calculation of motif enrichment and depletion in genomic windows

The local window enrichment plots in Fig. 4c and Fig. 4f were generated using the ‘enrich_in_sliding_windows’ function of SMEAGOL. This function creates windows tiling over the entire genome (for the figures here, non-overlapping windows of 500 bp were used) and tests whether the number of predicted binding sites for an RBP in each window is significantly higher / lower than the expected number based on a model in which binding sites for the RBP are uniformly distributed across the genome. P-values are calculated using a two-sided Fisher’s exact test and adjusted using the Benjamini-Hochberg procedure.

### Variant Effect Prediction

We downloaded information on 36,688 SARS-CoV-2 mutations from the GESS database (https://wan-bioinfo.shinyapps.io/GESS/) on September 14, 2021. We used the ‘variant_effect_on_sites’ function in SMEAGOL to select mutations that intersect with the predicted binding sites of 10 selected RBPs, and calculate the PWM match score of each predicted binding site with and without the variant. We selected variants that reduce the score of a binding site to less than 0.5 as site-disrupting variants.

## Supporting information

Supplementary Data 1: Strains and genotypes included in this study.

Supplementary Data 2: Locations of predicted RBP binding sites (sequences that match the PWM) on all viral genomes for all PWMs.

Supplementary Data 5: List of experimentally validated RBP effectors for SARS-CoV-2.

Supplementary Data 6: Selected mutations in the SARS-CoV-2 genome.

Supplementary Data 3: Enrichment / depletion results for all viral genomes and PWMs.

Supplementary Data 4: Enrichment / depletion results for all UTR sequences and PWMs.

## Data Availability

All results generated in this study are available in the supplementary materials. Accession numbers for external datasets used in this study are given in Supplementary Data 1. Our curated lists of PWMs (both full and representative sets) are available at https://github.com/gruber-sciencelab/VirusHostInteractionAtlas and the PWMs can also be loaded using SMEAGOL. Figures 2a, 2b, 3, 4b and 4e are based on raw data in Supplementary Data 3. Figure 4a is based on raw data in Supplementary Data 4. Figures 4c and 4f are based on raw data in Supplementary Data 2.

## Code Availability

SMEAGOL is available at https://github.com/gruber-sciencelab/SMEAGOL under an MIT open-source license. Our scripts and notebooks using SMEAGOL for the analysis of viral genomes and UTR sequences are available at https://github.com/gruber-sciencelab/VirusHostInteractionAtlas, also under an MIT open-source license.

## Acknowledgements

The authors thank Mihaela Zavolan for many constructive discussions and ideas that greatly improved this work. The authors acknowledge the use of de.NBI cloud and the support by the High Performance and Cloud Computing Group at the Zentrum für Datenverarbeitung of the University of Tübingen and the Federal Ministry of Education and Research (BMBF) through grant no 031 A535A, whereas special thanks go to Maximilian Hanussek and Jens Krüger for their help and great support with the cloud computing environment.

## Author Contributions

A.L. developed SMEAGOL with contributions from A.J.G. and M.G.F.. A.L. created Fig. 1, M.G.F. curated the viral genomes and created Figs. 2 and 3 based on data analyses performed by A.L., A.J.G. and M.G.F.. A.L. performed the SARS-CoV-2 analysis and A.J.G. the HCV analysis. All authors wrote and approved the manuscript.

## Supplementary Data Legends

Supplementary Data 1: Strains and genotypes included in this study.

Supplementary Data 3: Enrichment / depletion results for all viral genomes and PWMs

Supplementary Data 4: Enrichment / depletion results for all UTR sequences and PWMs.

Supplementary Data 5: List of experimentally validated RBP effectors for SARS-CoV-2.

Supplementary Data 6: Selected mutations in the SARS-CoV-2 genome.

## References

1. Baltimore, D. Expression of animal virus genomes. Bacteriol. Rev. 35, 235–241 (1971).

2. Flynn, R. A. et al. Discovery and functional interrogation of SARS-CoV-2 RNA-host protein interactions. Cell 184, 2394–2411.e16 (2021).

3. Schmidt, N. et al. The SARS-CoV-2 RNA-protein interactome in infected human cells. Nat Microbiol 6, 339–353 (2021).

4. Maranon, D. G., Anderson, J. R., Maranon, A. G. & Wilusz, J. The interface between coronaviruses and host cell RNA biology: Novel potential insights for future therapeutic intervention. Wiley Interdiscip. Rev. RNA doi:10.1002/wrna.1614.

5. Barnhart, M. D., Moon, S. L., Emch, A. W., Wilusz, C. J. & Wilusz, J. Changes in Cellular mRNA Stability, Splicing, and Polyadenylation through HuR Protein Sequestration by a Cytoplasmic RNA Virus. Cell Rep. 5, 909–917 (2013).

6. Cook, K. B., Kazan, H., Zuberi, K., Morris, Q. & Hughes, T. R. RBPDB: a database of RNA-binding specificities. Nucleic Acids Res. 39, D301–8 (2011).

7. Giudice, G., Sánchez-Cabo, F., Torroja, C. & Lara-Pezzi, E. ATtRACT-a database of RNA-binding proteins and associated motifs. Database 2016, (2016).

8. Van Nostrand, E. L. et al. A large-scale binding and functional map of human RNA-binding proteins. Nature 583, 711–719 (2020).

9. Villa, E. C., Maruyama, S. R., de Miranda-Santos, I. K. F., Palacios, G. & Ladner, J. T. Complete Coding Genome Sequence for Mogiana Tick Virus, a Jingmenvirus Isolated from Ticks in Brazil. Genome Announc. 5, (2017).

10. Viktorovskaya, O. V., Greco, T. M., Cristea, I. M. & Thompson, S. R. Identification of RNA Binding Proteins Associated with Dengue Virus RNA in Infected Cells Reveals Temporally Distinct Host Factor Requirements. PLoS Negl. Trop. Dis. 10, (2016).

11. Phillips, S. L., Soderblom, E. J., Bradrick, S. S. & Garcia-Blanco, M. A. Identification of Proteins Bound to Dengue Viral RNA In Vivo Reveals New Host Proteins Important for Virus Replication. MBio 7, (2016).

12. Sanford, T. J., Mears, H. V., Fajardo, T., Locker, N. & Sweeney, T. R. Circularization of flavivirus genomic RNA inhibits de novo translation initiation. Nucleic Acids Res. 47, 9789–9802 (2019).

13. Goldstrohm, A. C., Hall, T. M. T. & McKenney, K. M. Post-transcriptional Regulatory Functions of Mammalian Pumilio Proteins. Trends Genet. 34, 972–990 (2018).

14. Liu, Y. et al. Structures and Functions of the 3’ Untranslated Regions of Positive-Sense Single-Stranded RNA Viruses Infecting Humans and Animals. Front. Cell. Infect. Microbiol. 10, (2020).

15. Li, Z. & Nagy, P. D. Diverse roles of host RNA binding proteins in RNA virus replication. RNA Biology vol. 8 305–315 (2011).

16. Spångberg, K., Wiklund, L. & Schwartz, S. HuR, a protein implicated in oncogene and growth factor mRNA decay, binds to the 3’ ends of hepatitis C virus RNA of both polarities. Virology 274, 378–390 (2000).

17. Harris, D., Zhang, Z., Chaubey, B. & Pandey, V. N. Identification of cellular factors associated with the 3’-nontranslated region of the hepatitis C virus genome. Mol. Cell. Proteomics 5, 1006–1018 (2006).

18. Fan, B. et al. A Human Proteome Microarray Identifies that the Heterogeneous Nuclear Ribonucleoprotein K (hnRNP K) Recognizes the 5’ Terminal Sequence of the Hepatitis C Virus RNA. Mol. Cell. Proteomics 13, 84–92 (2014).

19. Korf, M., Jarczak, D., Beger, C., Manns, M. P. & Krüger, M. Inhibition of hepatitis C virus translation and subgenomic replication by siRNAs directed against highly conserved HCV sequence and cellular HCV cofactors. J. Hepatol. 43, 225–234 (2005).

20. Randall, G. et al. Cellular cofactors affecting hepatitis C virus infection and replication. Proc. Natl. Acad. Sci. U. S. A. 104, 12884–12889 (2007).

21. Rivas-Aravena, A. et al. The Elav-like protein HuR exerts translational control of viral internal ribosome entry sites. Virology 392, 178–185 (2009).

22. Shwetha, S. et al. HuR Displaces Polypyrimidine Tract Binding Protein To Facilitate La Binding to the 3’ Untranslated Region and Enhances Hepatitis C Virus Replication. J. Virol. 89, 11356–11371 (2015).

23. Chung, R. T. & Kaplan, L. M. Heterogeneous nuclear ribonucleoprotein I (hnRNP-I/PTB) selectively binds the conserved 3’ terminus of hepatitis C viral RNA. Biochem. Biophys. Res. Commun. 254, 351–362 (1999).

24. Gontarek, R. R. et al. hnRNP C and polypyrimidine tract-binding protein specifically interact with the pyrimidine-rich region within the 3’NTR of the HCV RNA genome. Nucleic Acids Res. 27, 1457–1463 (1999).

25. Ito, T. & Lai, M. M. An internal polypyrimidine-tract-binding protein-binding site in the hepatitis C virus RNA attenuates translation, which is relieved by the 3’-untranslated sequence. Virology 254, 288–296 (1999).

26. Chang, K.-S. & Luo, G. The polypyrimidine tract-binding protein (PTB) is required for efficient replication of hepatitis C virus (HCV) RNA. Virus Res. 115, 1–8 (2005).

27. Tsuchihara, K. et al. Specific interaction of polypyrimidine tract-binding protein with the extreme 3’-terminal structure of the hepatitis C virus genome, the 3’X. J. Virol. 71, 6720–6726 (1997).

28. Ali, N. & Siddiqui, A. Interaction of polypyrimidine tract-binding protein with the 5’ noncoding region of the hepatitis C virus RNA genome and its functional requirement in internal initiation of translation. J. Virol. 69, 6367–6375 (1995).

29. Beales, L. P., Rowlands, D. J. & Holzenburg, A. The internal ribosome entry site (IRES) of hepatitis C virus visualized by electron microscopy. RNA 7, 661–670 (2001).

30. Nomoto, A., Tsukiyama-Kohara, K. & Kohara, M. Mechanism of translation initiation on hepatitis C virus RNA. Princess Takamatsu Symp. 25, 111–119 (1995).

31. Zhang, J. et al. Down-regulation of viral replication by adenoviral-mediated expression of siRNA against cellular cofactors for hepatitis C virus. Virology 320, 135–143 (2004).

32. Domitrovich, A. M., Diebel, K. W., Ali, N., Sarker, S. & Siddiqui, A. Role of La autoantigen and polypyrimidine tract-binding protein in HCV replication. Virology 335, 72–86 (2005).

33. Huang, P. & Lai, M. M. Polypyrimidine tract-binding protein binds to the complementary strand of the mouse hepatitis virus 3’ untranslated region, thereby altering RNA conformation. J. Virol. 73, 9110–9116 (1999).

34. Babic, I., Sharma, S. & Black, D. L. A role for polypyrimidine tract binding protein in the establishment of focal adhesions. Mol. Cell. Biol. 29, 5564–5577 (2009).

35. Görlach, M., Burd, C. G. & Dreyfuss, G. The determinants of RNA-binding specificity of the heterogeneous nuclear ribonucleoprotein C proteins. J. Biol. Chem. 269, 23074–23078 (1994).

36. Wan, L., Kim, J. K., Pollard, V. W. & Dreyfuss, G. Mutational definition of RNA-binding and protein-protein interaction domains of heterogeneous nuclear RNP C1. J. Biol. Chem. 276, 7681–7688 (2000).

37. König, J. et al. iCLIP reveals the function of hnRNP particles in splicing at individual nucleotide resolution. Nat. Struct. Mol. Biol. 17, 909–915 (2010).

38. Li, Y., Yamane, D., Masaki, T. & Lemon, S. M. The yin and yang of hepatitis C: synthesis and decay of hepatitis C virus RNA. Nat. Rev. Microbiol. 13, 544–558 (2015).

39. Samuel, C. E. The eIF-2 alpha protein kinases, regulators of translation in eukaryotes from yeasts to humans. J. Biol. Chem. 268, 7603–7606 (1993).

40. Buchan, J. R. & Parker, R. Eukaryotic stress granules: the ins and outs of translation. Mol. Cell 36, 932–941 (2009).

41. Garaigorta, U., Heim, M. H., Boyd, B., Wieland, S. & Chisari, F. V. Hepatitis C virus (HCV) induces formation of stress granules whose proteins regulate HCV RNA replication and virus assembly and egress. J. Virol. 86, 11043–11056 (2012).

42. Dixit, U. et al. FUSE Binding Protein 1 Facilitates Persistent Hepatitis C Virus Replication in Hepatoma Cells by Regulating Tumor Suppressor p53. J. Virol. 89, 7905–7921 (2015).

43. Chatel-Chaix, L. et al. A host YB-1 ribonucleoprotein complex is hijacked by hepatitis C virus for the control of NS3-dependent particle production. J. Virol. 87, 11704–11720 (2013).

44. Diosa-Toro, M., Prasanth, K. R., Bradrick, S. S. & Garcia Blanco, M. A. Role of RNA-binding proteins during the late stages of Flavivirus replication cycle. Virol. J. 17, 1–14 (2020).

45. Kawaguchi, A., Matsumoto, K. & Nagata, K. YB-1 functions as a porter to lead influenza virus ribonucleoprotein complexes to microtubules. J. Virol. 86, (2012).

46. Jung, Y.-M. et al. Investigation of function and regulation of the YB-1 cellular factor in HIV replication. BMB Rep. 51, 290 (2018).

47. Lee, S. et al. The SARS-CoV-2 RNA interactome. Mol. Cell 81, 2838–2850.e6 (2021).

48. Kamel, W. et al. Global analysis of protein-RNA interactions in SARS-CoV-2-infected cells reveals key regulators of infection. Mol. Cell 81, 2851–2867.e7 (2021).

49. Wei, J. et al. Genome-wide CRISPR Screens Reveal Host Factors Critical for SARS-CoV-2 Infection. Cell 184, 76–91.e13 (2021).

50. Fang, S. et al. GESS: a database of global evaluation of SARS-CoV-2/hCoV-19 sequences. Nucleic Acids Res. 49, D706–D714 (2021).

51. Resende, P. C. et al. A Potential SARS-CoV-2 Variant of Interest (VOI) Harboring Mutation E484K in the Spike Protein Was Identified within Lineage B.1.1.33 Circulating in Brazil. Viruses 13, (2021).

52. Heinz, S. et al. Simple combinations of lineage-determining transcription factors prime cis-regulatory elements required for macrophage and B cell identities. Mol. Cell 38, 576–589 (2010).

53. Korhonen, J., Martinmäki, P., Pizzi, C., Rastas, P. & Ukkonen, E. MOODS: fast search for position weight matrix matches in DNA sequences. Bioinformatics 25, 3181–3182 (2009).

54. Tan, G. & Lenhard, B. TFBSTools: an R/bioconductor package for transcription factor binding site analysis. Bioinformatics 32, 1555–1556 (2016).

55. Wasserman, W. W. & Sandelin, A. Applied bioinformatics for the identification of regulatory elements. Nat. Rev. Genet. 5, 276–287 (2004).

56. motifmatchr. http://bioconductor.org/packages/release/bioc/html/motifmatchr.html.

57. Stasi, C., Silvestri, C. & Voller, F. Update on Hepatitis C Epidemiology: Unaware and Untreated Infected Population Could Be the Key to Elimination. SN Compr Clin Med 1–8 (2020).

58. Herdy, B. et al. The RNA-binding protein HuR/ELAVL1 regulates IFN-β mRNA abundance and the type I IFN response. Eur. J. Immunol. 45, 1500–1511 (2015).

59. Takeuchi, O. HuR keeps interferon-β mRNA stable. Eur. J. Immunol. 45, 1296–1299 (2015).

60. Ferrarini, M. G. et al. Genome-wide bioinformatic analyses predict key host and viral factors in SARS-CoV-2 pathogenesis. Commun Biol 4, 590 (2021).

61. Alipanahi, B., Delong, A., Weirauch, M. T. & Frey, B. J. Predicting the sequence specificities of DNA- and RNA-binding proteins by deep learning. Nat. Biotechnol. 33, 831–838 (2015).

62. Park, C. Y. et al. Genome-wide landscape of RNA-binding protein target site dysregulation reveals a major impact on psychiatric disorder risk. Nat. Genet. 53, 166–173 (2021).

63. Grønning, A. G. B. et al. DeepCLIP: predicting the effect of mutations on protein-RNA binding with deep learning. Nucleic Acids Res. 48, 7099–7118 (2020).

64. Shrikumar, A. et al. Technical Note on Transcription Factor Motif Discovery from Importance Scores (TF-MoDISco) version 0.5.6.5. arXiv [cs.LG] (2018).

65. Phadke, S., Macherla, S. & Scheuermann, R. H. Database and Analytical Resources for Viral Research Community. Reference Module in Life Sciences (2019).

66. Hulo, C. et al. ViralZone: a knowledge resource to understand virus diversity. Nucleic Acids Res. 39, D576–82 (2011).

67. Castro-Mondragon, J. A., Jaeger, S., Thieffry, D., Thomas-Chollier, M. & van Helden, J. RSAT matrix-clustering: dynamic exploration and redundancy reduction of transcription factor binding motif collections. Nucleic Acids Res. 45, e119–e119 (2017).

