## Supplementary Materials for "Investigating the human host - ssRNA virus interaction landscape using the SMEAGOL toolbox"

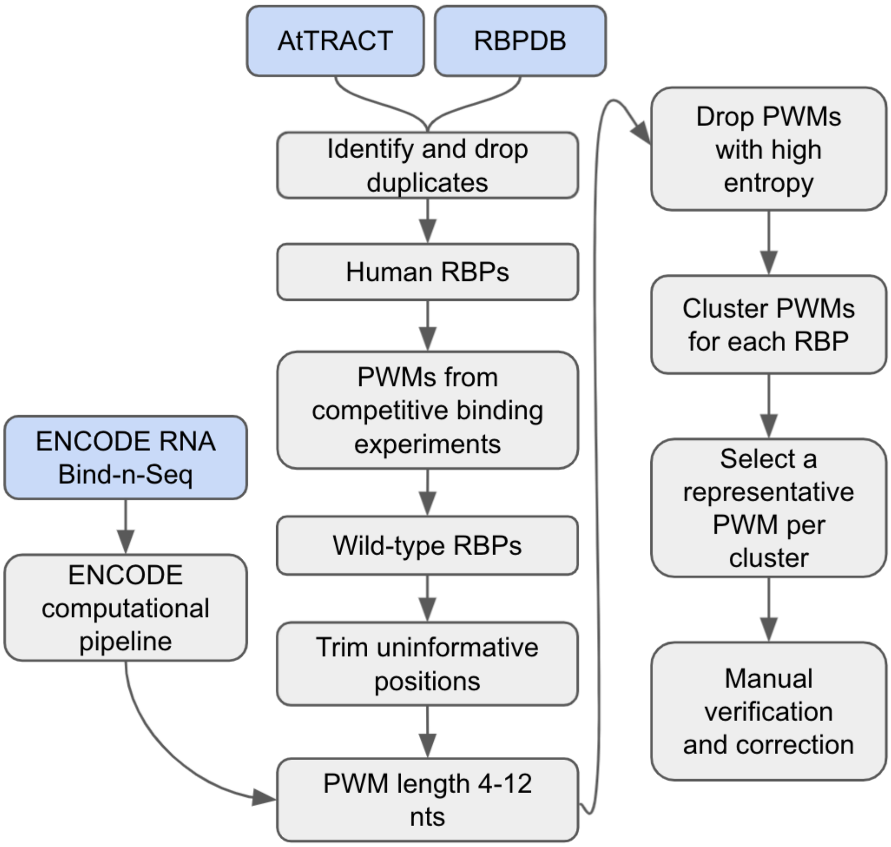


**Supplementary Figure 1:** Schematic of PWM curation strategy.

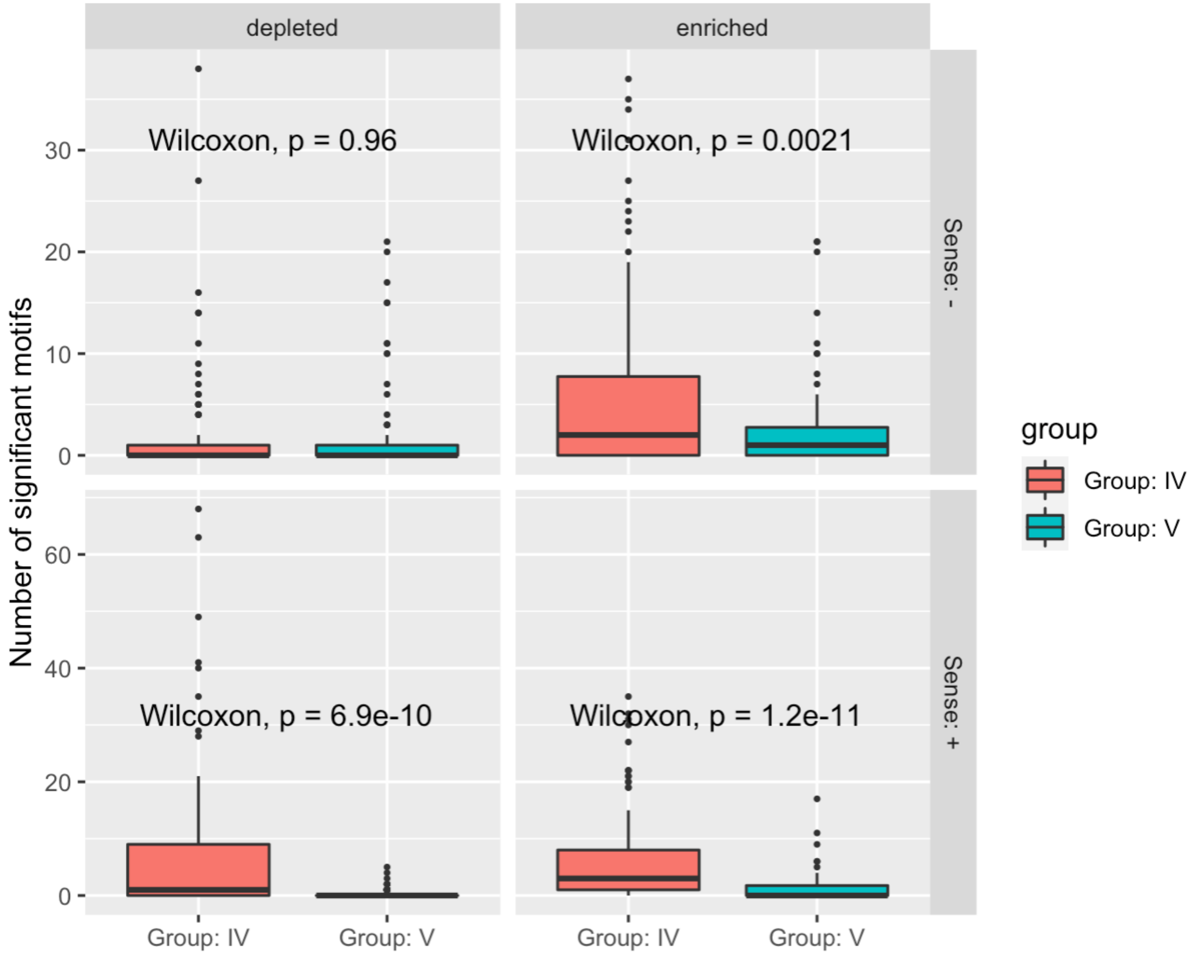


**Supplementary Figure 2**: Boxplots showing the number of motifs in our dataset that were enriched or depleted per genome in Group IV or Group V viruses, on the + and - sense sequences. Box plots are defined as follows: center line, median; box limits, upper and lower quartiles; whiskers, 1.5x interquartile range; points, outliers.

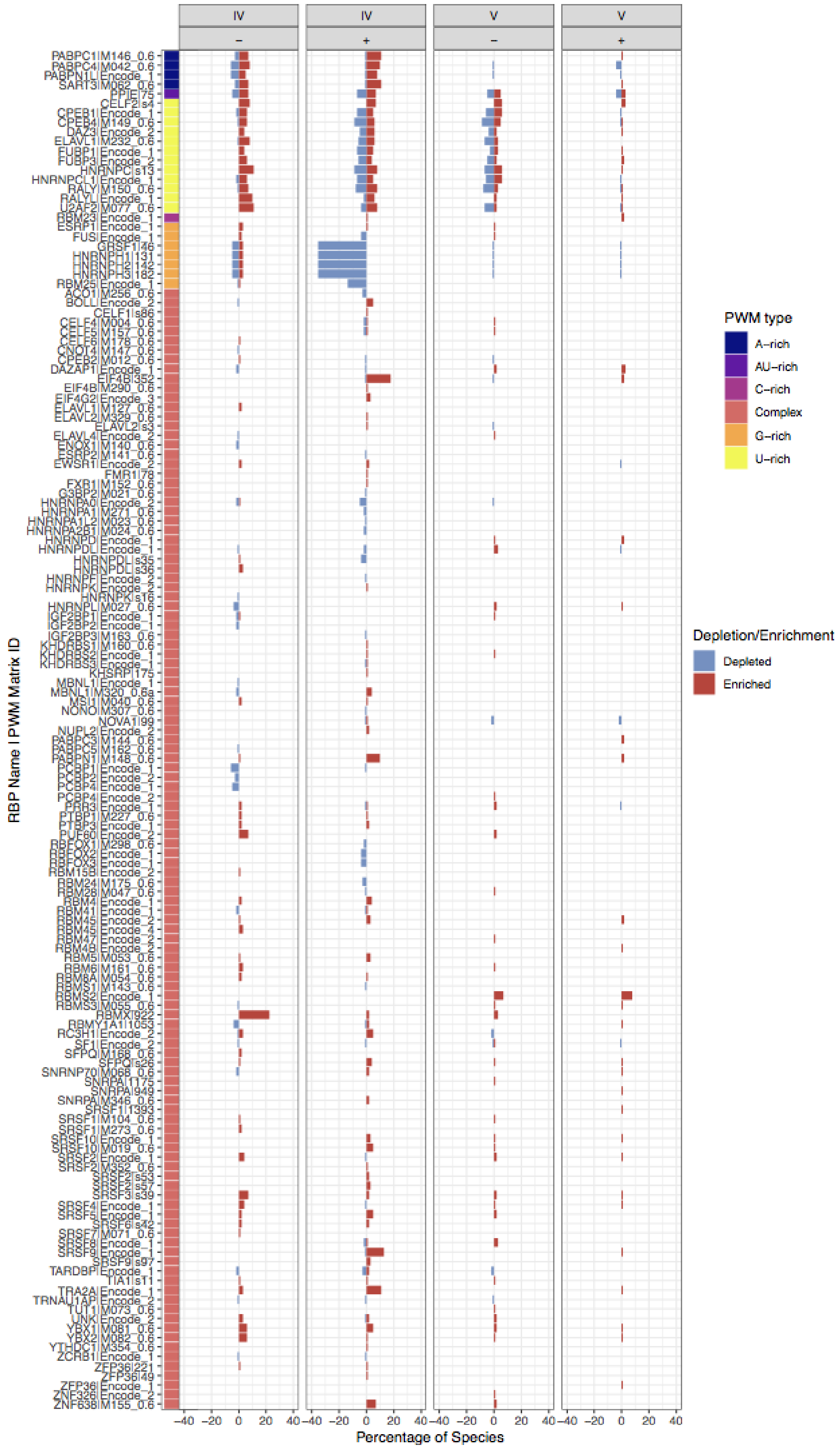


**Supplementary Figure 3:** Number of occurrences of enrichment (in red) and depletion (in blue) for all cluster-representative PWMs, per virus class (IV or V) and per strand, normalized per 100 genomes. PWM sequence bias is presented on the left side of the plot: while some PWMs have a more complex sequence, others are rich in single nucleotides.

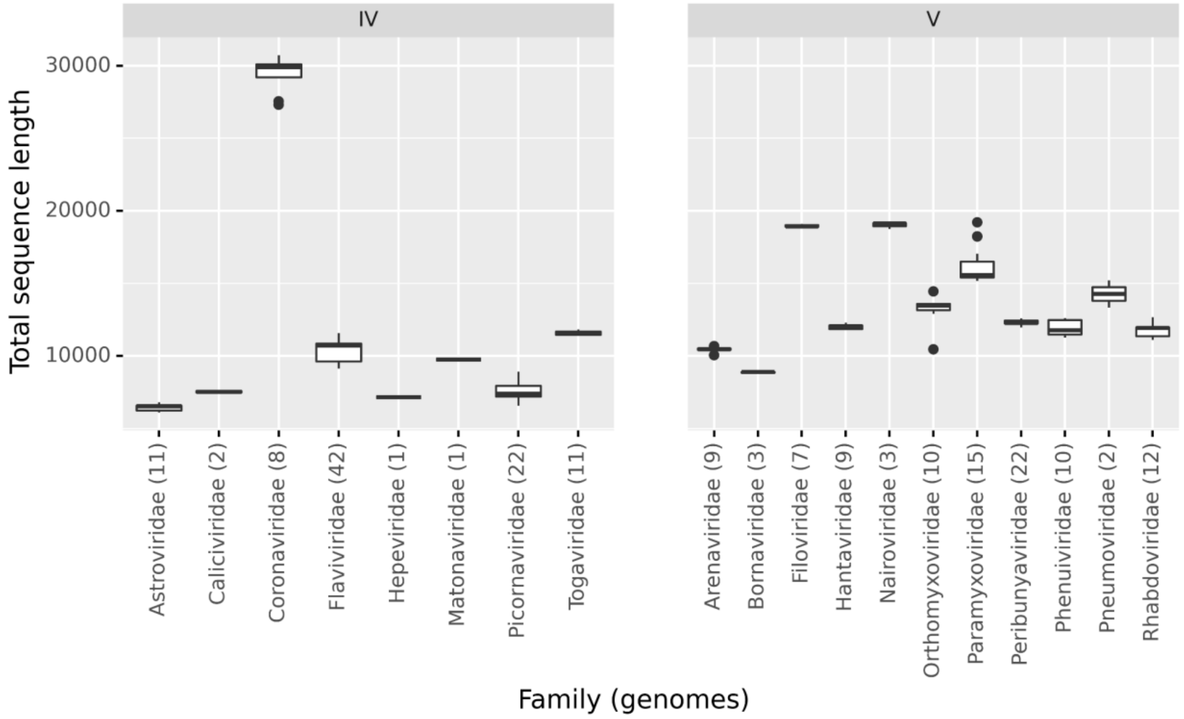


**Supplementary Figure 4:** Boxplots showing genome lengths in different viral families. The number of genomes for each family is given in parentheses. Box plots are defined as follows: center line, median; box limits, upper and lower quartiles; whiskers, 1.5x interquartile range; points, outliers.

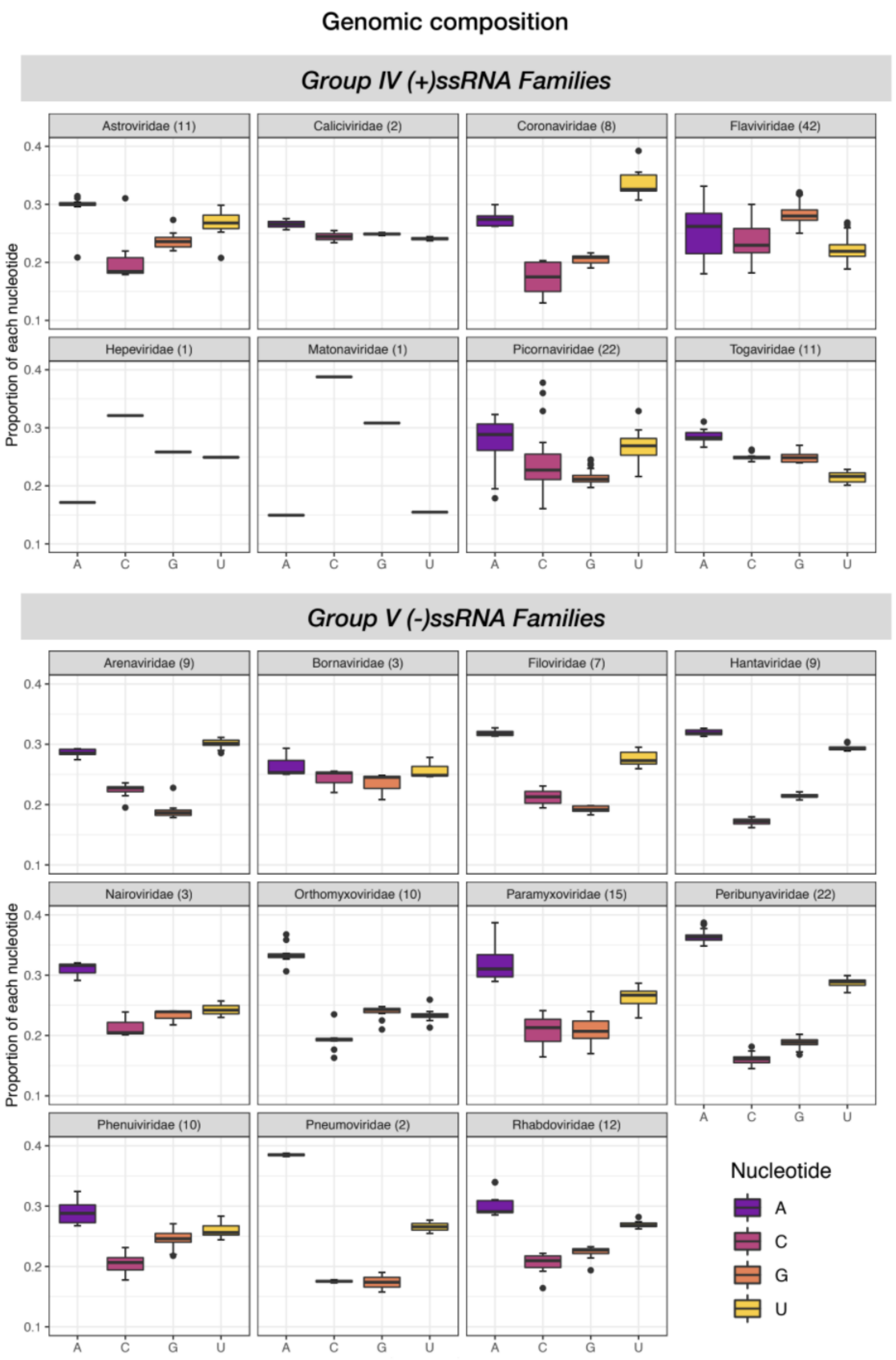


**Supplementary Figure 5:** Boxplots showing base composition of the (+) sense genome in different viral families. The number of genomes for each family is given in parentheses. Box plots are defined as follows: center line, median; box limits, upper and lower quartiles; whiskers, 1.5x interquartile range; points, outliers.

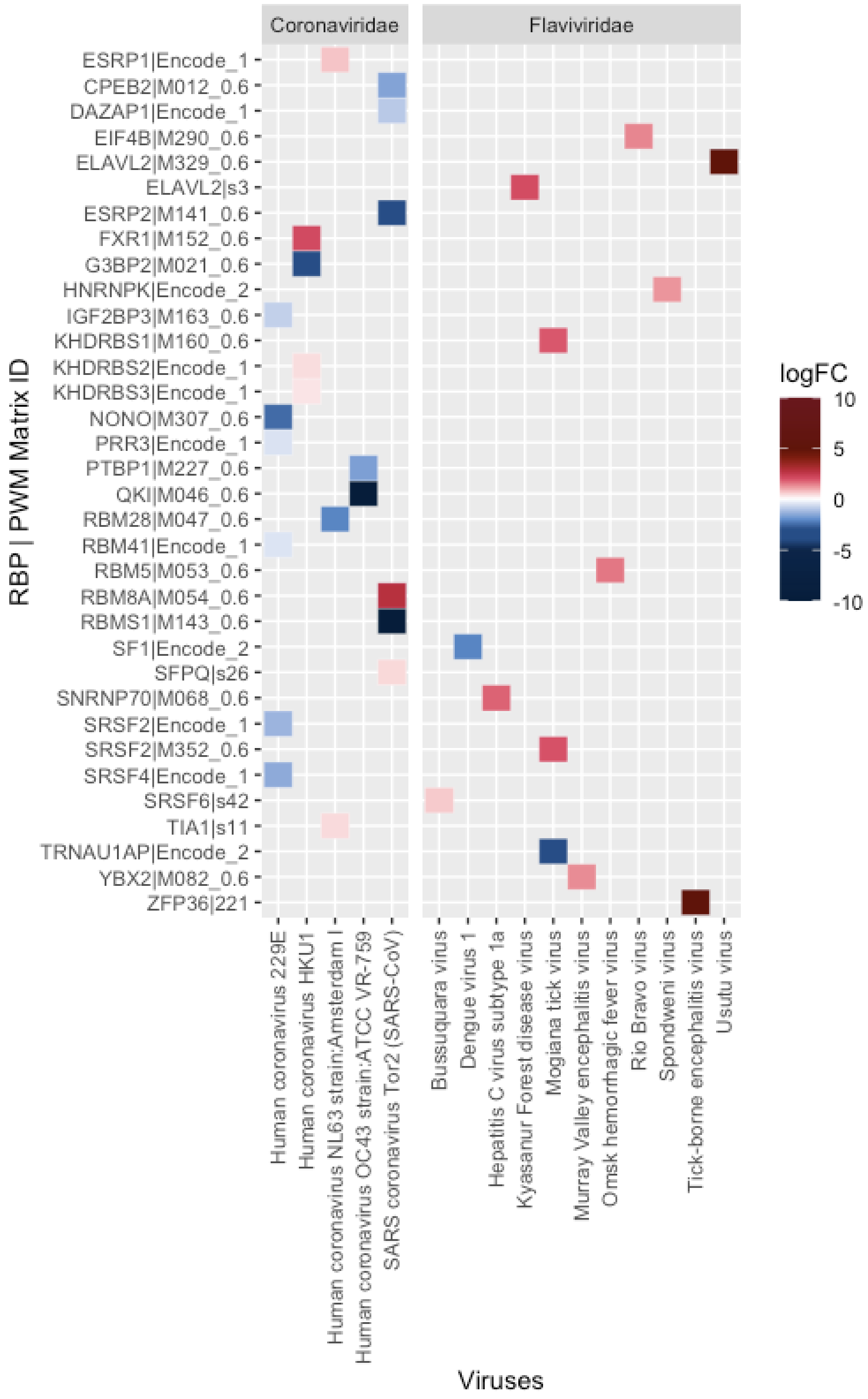


**Supplementary Figure 6.** Heatmap for species specific enrichment / depletion results within *Coronaviridae* and *Flaviviridae* families.

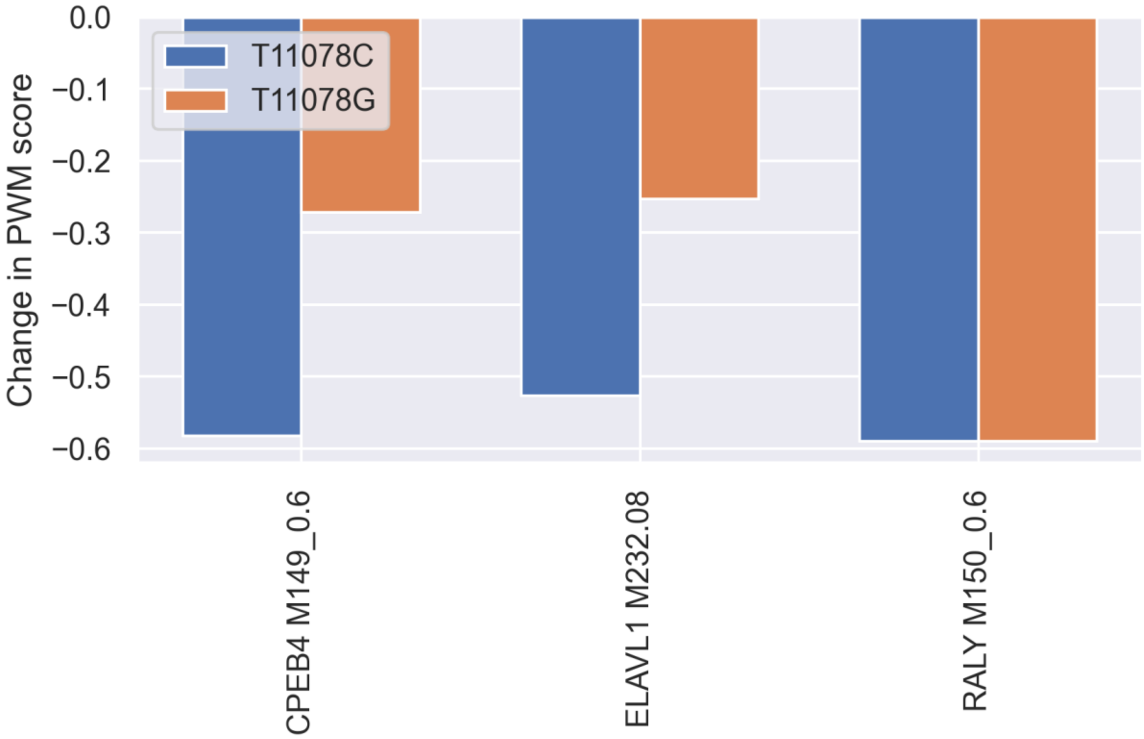


**Supplementary Figure 7:** Predicted effect of T>C and T>G mutations at position 11078 of the SARS-CoV-2 genome, on binding of three RBPs (CPEB4, ELAVL1 and RALY).

**a**

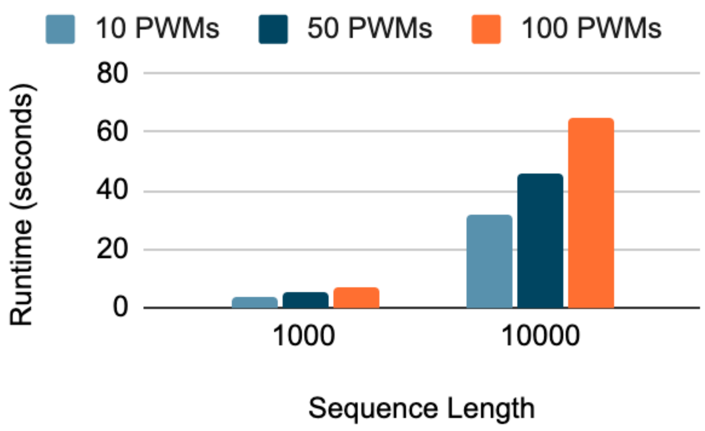


**b**

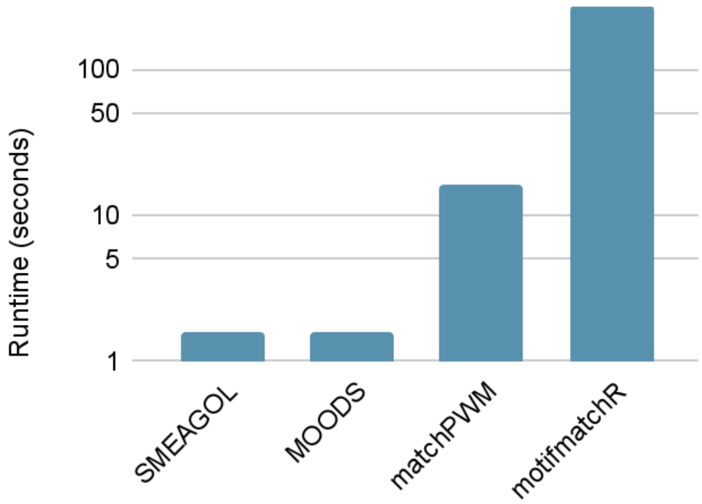


**Supplementary Figure 8:** a) Time in seconds to perform complete enrichment analysis using SMEAGOL for given numbers of PWMs, on a sequence of length 1’000 or 10’000 bases. b) Time in seconds to scan 100 sequences each of length 10’000 bases with 50 PWMs, using SMEAGOL or other PWM scanning softwares. Benchmarks were performed on a 14 cores node with 32 GB of memory.
